# Oncoprotein 18 is necessary for malignant cell proliferation in bladder cancer cells and serves as a G3-specific non-invasive diagnostic marker candidate in urinary RNA

**DOI:** 10.1101/2020.02.03.932947

**Authors:** Merle Hanke, Josephine Dubois, Ingo Kausch, Sonja Petkovic, Georg Sczakiel

**Author notes:** to whom correspondence should be addressed: Georg Sczakiel.

## Abstract

**Background:** Urine-based diagnostics indicated involvement of OP18 in bladder cancer. In cell culture models we investigated the role of oncoprotein 18 for malignant cell growth.

**Methods:** We analyzed 113 urine samples and investigated two human BCa cell lines as a dual model: RT-4 and ECV-304, which represented differentiated (G1) and poorly differentiated (G3) BCa. We designed specific siRNA for down-regulation of OP18 in both cell lines. Phenotypes were characterized by cell viability, proliferation, and expression of apoptosis-related genes. Besides, sensitivity to cisplatin treatment was evaluated.

**Results:** Analysis of urine samples from patients with urothelial BCa revealed a significant correlation of the RNA-ratio oncoprotein 18:uroplakin 1A with bladder cancer. High urinary ratios were mainly found in moderately to poorly differentiated tumors (grade G2-3) that were muscle invasive (stage T2-3), whereas samples from patients with more differentiated non-invasive BCa (G1) showed low OP18:UPK1A RNA ratios. Down-regulation of OP18 expression in ECV-304 shifted its phenotype towards G1 state. Further, OP18-directed siRNA induced apoptosis and increased chemo-sensitivity to cisplatin.

**Conclusions:** This study provides conclusive experimental evidence for the link between OP18-derived RNA as a diagnostic marker for molecular staging of BCa in non-invasive urine-based diagnostics and the patho-mechanistic role of OP18 suggesting this gene as a therapeutic target.

## 1 INTRODUCTION

The treatment of bladder cancer (BCa) depends on its stage. While non-muscle invasive forms of BCa can be removed by TUR-B of tumor tissue and its recurrence can be treated by immunotherapy with intra-vesicular delivery of attenuated *Bacillus Calmette–Guérin* (BCG) or intra-vesical chemotherapy, muscle-invasive tumor forms demand more aggressive strategies. Chemotherapy includes platinum-based drugs like cis-diamminedichloridoplatinum(II), (henceforth referred to as cisplatin), as one of the standard chemotherapeutic agents for the treatment of metastatic BCa [1, 2].

The efficacy of a multiplicity of chemotherapeutic agents including cisplatin is often substantially decreased since BCa tumors frequently develop a drug- or multiple drug-resistance (MDR) mechanisms [3, 4]. Drug-resistant cells show, amongst others, an over-expression of anti-apoptotic genes [4]. Hence, the identification of new molecular targets and alternative classes of drugs, including oligonucleotide-based medications [5, 6], is crucial for the improvement of survival rates of patients with advanced BCa. There is a high clinical interest in objective and more accurate ways of tumor classification that may replace tissue-based histopathological staging. Innovative diagnostic approaches are increasingly based on the non-invasive monitoring of BCa-specific tumor markers in urine. Promising markers for bladder cancer had been based on RNA such as microRNAs and also sequences of cellular mRNAs [7–9]. Further, we showed that analysis of the RNA composition in whole urine of BCa patients reveals specific and sensitive RNA-based tumor markers including ETS2 and uPA [10] as well as microRNAs [11].

In this study, we aimed to investigate whether differentially detectable RNAs in whole urine of BCa patients provide improved tumor markers *per se*. Further, we asked whether those RNA markers might display aberrant mRNA expression (of OP18) in malignant cells. We used two different BCa cell culture models to evaluate a possible involvement of OP18 gene expression in the tumorigenesis: The human cell lines ECV-304 [12–14] and RT-4 [15], representing well (G1) and poorly differentiated (G3) tumor states, respectively. To test whether model-based G1/G3-related phenotypes were in line with our results in liquid biopsies, we quantified RNA and protein levels of OP18 and performed several proliferation and cell viability assays under siRNA mediated suppression of OP18 mRNA. We also analyzed the influences of OP18 on apoptotic genes and chemo-sensitivity of G3 cells.

## 2 MATERIALS AND METHODS

### 2.1 Clinical samples and preparation of RNAs

This study was approved by the local ethical research committee in Lübeck. All urine samples were obtained with written informed consents of the participants. Tumor grading was determined by urinary bladder cystoscopy. In addition to biopsy, urine cytology was performed. All tumors identified were completely resected and classified pathologically according to the World Health Organization 1973 grading and staging system. For investigation of urinary OP18 and uroplakin 1A (UPK1A) RNA levels, spontaneously voided urine of 113 donors was collected: 61 BCa patients (male:female, 3:1; G1 pTa, n = 13; G2+G3 pTa+pTis, n = 19; G2+G3 pT1, n = 12; G2+G3 pT2+pT3, n = 17; median age, 71 years), 37 healthy volunteers (male:female, 2:1; median age, 71 years), and 15 patients with infections of the urinary tract (male:female, 1:7; median age, 55 years). Urine samples were stabilized immediately as described recently [11].

Total RNA from cells was prepared using the RNeasy Mini kit (Qiagen, Hilden, Germany), and urinary RNA was isolated using the RNeasy Midi Kit (Qiagen, Hilden, Germany), except for the lysis buffer described above which was used instead of buffer “RLT”. RNA was eluted twice with 160 μl H_2_O and then lyophilized. Pellets were resolved in 20 μl H_2_O, and the quality of RNA was assessed by agarose gel electrophoresis (using 1 μg/ml ethidium bromide). Urinary RNA extract (10 μl) or 400 ng total RNA from cells were used for cDNA synthesis and minus reverse transcriptase (non-RT) reaction.

### 2.2 cDNA synthesis

Reverse transcription was performed in a total volume of 20 μl with RNA extract (10 μl), and 300 ng random hexamer primer (Invitrogen, Paisley, UK) following the manufacturer’s instructions for SuperScript III™ driven reverse transcription (Invitrogen, Paisley, UK), despite a little increase of time and temperature for the denaturation 75°C and 10 min.

### 2.3 Quantitative PCR (qPCR) and data analysis

Primers and TaqMan^®^ probes were designed using Primer Express^®^ software version 2.0 (Applied Biosystems, Darmstadt, Germany) or Primer3 software (Steve Rozen, Whitehead Institute for Biomedical Research, Cambridge, UK) and were purchased from Metabion (Martinsried, Germany) and Eurogentec (Seraing, Belgium), respectively. Primer sequences were checked for homology using the Blast software (www.ncbi.nlm.nih.gov/BLAST). Amplicon characteristics and software information are listed in the S1 Table.

All reactions were performed with the qPCR Core Kit (Eurogentec, Seraing, Belgium) in a total reaction volume of 10 μl in 384-well plates. A non-template control (nuclease-free H_2_O) was included for each amplicon to exclude contamination in every qPCR run. Each qPCR reaction was performed in triplicate. Quantitative PCR was carried out in a 7900HT thermal cycler (Applied Biosystems, Darmstadt, Germany): initial denaturation at 95°C for 10 min, followed by 50 cycles at 95°C for 15 sec and 60°C for 60 sec. PCR products were purified using the MinElute PCR Purification Kit (Qiagen, Hilden, Germany). Six serial 10-fold dilutions (10^1^–10^6^ copy numbers/reaction) were prepared in 10 mmol/l Tris/HCl (pH 8.0), 10 ng/ml polyinosinic acid potassium salt to generate standard curves. Data analysis was performed via the SDS 2.1 software (Applied Biosystems, Darmstadt, Germany) and the *threshold cycle (Ct)* values of amplified targets were transformed into absolute RNA copy numbers using the standard curves.

### 2.4 Cell culture

The human urinary BCa cell line ECV-304 was cultivated in Medium 199 (with HEPES buffer + Earle’s salts) containing 10% (vol/vol) fetal calf serum (FCS Gold). ECV-304 was originally established from an invasive, G3 BCa of an 82 years old Swedish female patient with a mutant p53 in 1970 and is a defined derivative of T-24 [12–14] (from DSMZ, ACC-310, cell identity was confirmed by DNA profiling in September 2017 by the DSMZ). RT-4 [15] (from DSMZ, ACC-412, purchased in March 2018) was cultivated in RPMI 1640 supplemented with 10% (vol/vol) fetal calf serum and used as an *in vitro* model for differentiated G1 BCa. Both cell lines grew without antibiotics in culture medium at 37°C and 5% CO2 in a humidified incubator. All culture media and supplements were obtained from PAA Laboratories GmbH (Pasching, Austria). Control for Mycoplasma contamination was done using Venor^®^GeM Mycoplasma Detection Kit (Minerva Biolabs, Berlin Deutschland) according to manufacturer’s protocol.

### 2.5 Design and validation of siRNAs

Two small interfering RNAs targeting OP18 mRNA were designed in silico according to a systematic computational analysis of local target mRNA structures as described previously [16]. An extensive BLAST search indicated that both siRNA sequences were target-specific. Nucleotide sequences of the effective siRNA and a scrambled control siRNA without homology to human sequences are shown in the S2 Table. For the annealing of RNA strands (from IBA Goettingen, Germany), 20 μM of the sense and antisense strand, respectively, was denaturated in buffer (50 mmol/l potassium acetate, 1 mmol/l magnesium acetate, 15 mmol/l HEPES (pH 7.4)) at 90°C for 2 min followed by an annealing step at 37°C for 1 h. For transfection of siRNA, cells were seeded into tissue culture plates (12-well: 5 × 10^4^ ECV-304 cells or 8 × 10^4^ RT-4 cells; 96-well plates: 3 × 10^3^ ECV-304 cells). After 24 h cells were transfected for 4 h at 37°C with 30 nM of siRNA using Lipofectamine 2000 in OPTI-MEM I according to the manufacturer’s instructions (Invitrogen, Paisley, UK).

### 2.6 Phenotypic characterization and cell proliferation

For analysis of cell proliferation post-transfection with siRNA (12-well plate, 8 days, or in 96-well for 4 days) 3-5 × 10^3^ ECV-304 cells or 1 × 10^4^ RT-4 cells were seeded in 12-well plates with 1 ml of culture medium or 0.1 ml in 96-well plates, respectively. After 18-24 h, cells were transfected with siRNAs as described above. Every 2 days, 500 μl for 12-well (and 50 μl for 96-well) plates of culture medium was replaced. At time points of measurement, cells were harvested, and trypan blue-negative cells were counted using a Neubauer hemocytometer (Sigma Aldrich, Steinheim, Germany).

At day 1, 2, 3 and 4, cell viability was determined using a tetrazolium salt-derived (3-(4,5-dimethylthiazol-2-yl)-2,5-diphenyltetrazolium bromide (MTS)) colorimetric assay. Cell culture medium was replaced by pH indicator-free culture medium containing 0.32 mg/ml MTS (Promega, Mannheim, Germany) and 0.0073 mg/ml phenazine methosulfate (Sigma-Aldrich). Cells were cultivated at 37 °C for 2.5 h and A_490_ was determined by a Tecan Sunrise ELISA reader (Tecan Deutschland GmbH, Crailsheim, Germany).

Chemo-sensitivity of ECV-304 cells after siRNA-mediated suppression of OP18 was determined by treatment with cisplatin (Cisplatin-Teva^®^, Teva Pharma AG, Aesch, Switzerland) at 0, 1, 3, 6, 9, 12 μg/ml cisplatin over a period of 24 h starting at day 2 after transfection. Cell viability was determined using the MTS assay.

### 2.7 Detection of apoptosis

ECV-304 cells were seeded (3 × 10^3^ cells, white-bottom 96-well plates) and transfected after 24 h. Caspase 3/7 activity, was determined at day 3 and 4 post-transfection using the Caspase-Glo 3/7 Assay (Promega, Mannheim, Germany). The emerging fluorescence was detected (485_Ex_/527_Em_; Labsystems Fluoroscan Ascent, Helsinki, Finland), caspase 3/7 substrate was added, and bioluminescence detected after incubation at room temperature for approximately 2.5 h and normalized to cell viability.

### 2.8 Western blot analysis

ECV-304 cells were seeded in 12-well plates (5 × 10^4^ cells/well) and transfected after 24 h using the protocol described above. At 0, 1, 2, 3, 4 days post-transfection, cells were trypsinized (0.05% trypsin/0.02% EDTA in 1x PBS for 5 min at 37°C), washed (PBS 1x ice-cold) and lysed (buffer containing 20% glycerol, 2% SDS, 125 mM Tris/HCl (pH 6.8), 5% beta-mercaptoethanol, and 0.02 % (w/v) bromphenolblue). After denaturation (95°C for 5 min), samples were centrifuged (20 000 g for 1 min) and loaded onto a 16% SDS-polyacrylamide gel. Protein amounts of OP18 and beta-actin were quantified using a primary stathmin polyclonal antibody (1:1000; Cell Signaling Technology, NEB GmbH, Frankfurt/Main, Germany) and a polyclonal beta-actin antibody (1:1000; Abcam, Acris Antibodies, Hiddenhausen, Germany).Primary antibodies were detected by a secondary anti-rabbit IgG antibody conjugated with horseradish peroxidase (Dako, Glostrup, Denmark) and visualized via chemiluminescence (Pierce, Thermo Scientific, Karlsruhe, Germany).

## 3 RESULTS

### 3.1 Increased urinary OP18:UPK1A RNA-ratios are associated with invasive BCa

Total RNA was prepared from whole urine samples of healthy donors, patients with urinary tract infections, and patients with BCa. Analysis of urinary RNA of revealed an RNA signal ratio OP18:UPK1A which is significantly (*p* < 0.001) increased in patients with poorly differentiated (G3) BCa as well as invasive BCa (≥ pT2) (Fig. 1 A).

**Fig 1.**
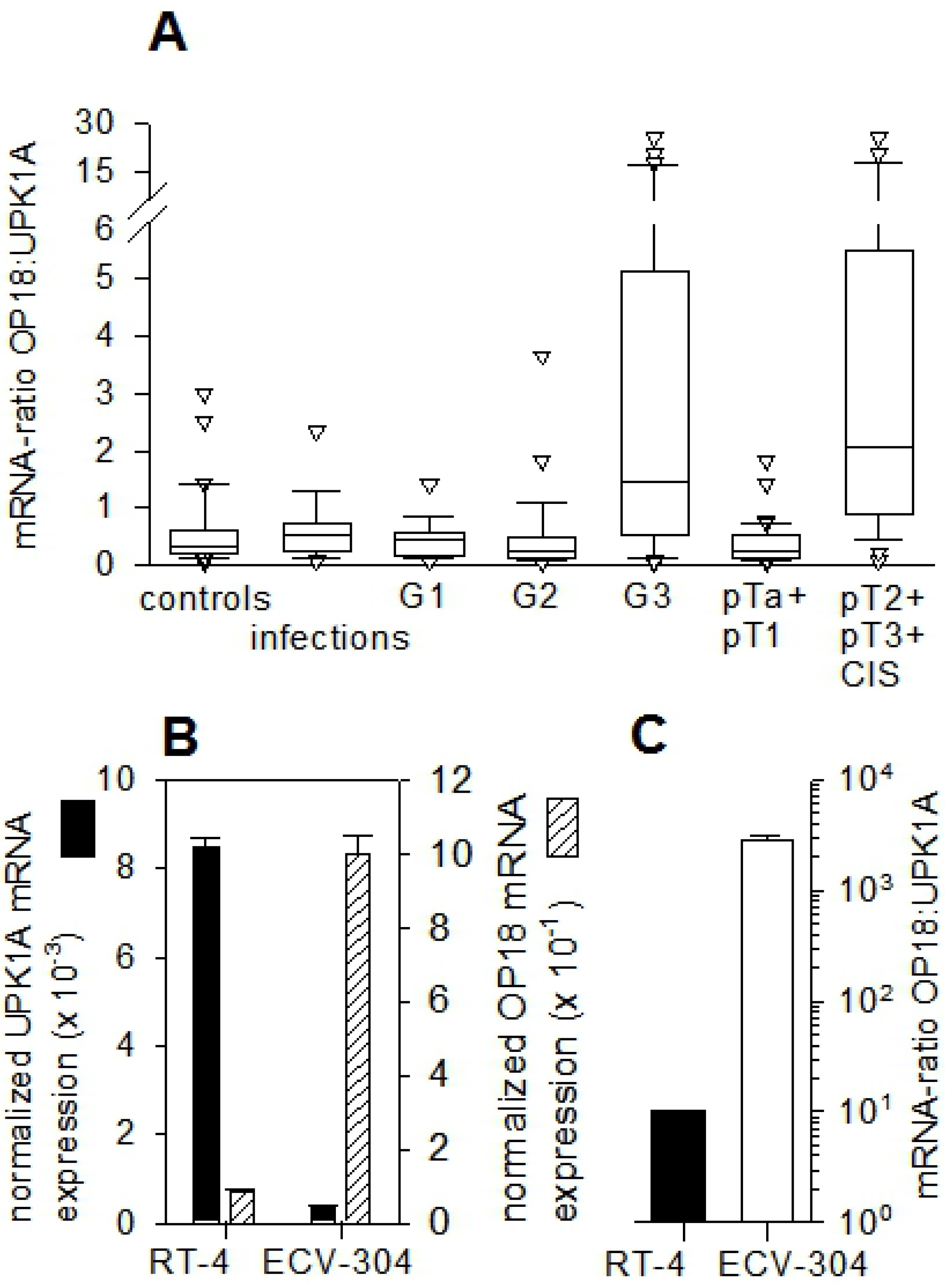
Urinary levels of the mRNA-based tumor marker OP18:UPK1A. **(A)**, Box-Plot of the urinary mRNA ratio OP18:UPK1A. The qRT-PCR-based detection of OP18- and UPK1A mRNA was performed in triplicate. Donors were of different health status, including healthy individuals and individuals suffering from infections of the urinary tract or BCa as stratified according to grade (G1, G2 or G3) and stage, respectively (see Materials and Methods for details). Upper and lower limits of boxes and lines across boxes indicate the 75^th^ and 25^th^ percentiles and median, respectively. Error bars indicate the 90^th^ and 10^th^ percentiles. White triangles indicate outlying data points. **(B)**, UPK1A- and OP18 mRNA expression in human BCa cell lines. Total RNA was prepared in triplicate from ECV-304 and RT-4 cells in the exponential phase of growth. Absolute mRNA copy numbers of OP18, UPK1A, and RPLP0 (internal reference mRNA) were determined in triplicate by qRT-PCR using standard curves. Columns represent mean values ± standard deviation of UPK1A and OP18 mRNA copies after normalization to RPLP0 mRNA levels. **(C)**, the abundance of the mRNA-ratio OP18:UPK1A in RT-4 and ECV-304 (calculated from single mRNA expression profiles as presented in Fig. 1 B). To compare OP18 to UPK1A-mRNA ratios of three or more patient groups, the Kruskal-Wallis test (H-test) and for two different patient groups the unpaired Mann-Whitney-U test was used. For all statistical tests, two-sided P values ≤ 0.05 were considered as statistically significant.

While UPK1A-specific RNA sequences were less abundant in urine samples from G3 BCa patients when compared to G2 BCa patients, the high level of urinary OP18-specific RNA sequences increased with tumor invasiveness, thereby representing the determining factor for an increased OP18:UPK1A mRNA-ratio (Fig. 1).

Besides the potential suitability of OP18-derived RNA as a urinary marker for molecular staging, this observation indicated an involvement of OP18 gene expression in the tumorigenesis of BCa. To test whether model-based G1/G3-related RNA levels of OP18 and UPK1A were compatible with those in liquid biopsies, total RNA was prepared from both cell lines: The BCa-derived human cell lines RT-4 and ECV-304, representing well (G1) and poorly differentiated (G3) tumor states, respectively. Absolute copy numbers of OP18 and UPK1A as well as RPLP0 RNA (60S acidic ribosomal protein P0, serving as internal reference mRNA) were determined via qRT-PCR. This test revealed matching RNA levels of OP18 and UPK1A, respectively, and suggested the validity of this cell-based system to study the role of OP18 for malignant cell proliferation of bladder cancer (Fig. 1 B and C).

### 3.2 Validation of siRNA tools against OP18 translation

The siRNA (sequences in S2 Table) was tested for suppression of OP18 expression in 5 × 10^4^ ECV-304 cells or 8 × 10^4^ RT-4 cells. The concentration dependency of siRNA-mediated suppression of OP18 at the transcriptional level showed an IC_50_ value of the most effective OP18-directed siRNA in ECV-304 cells in the range of 100 pM. Time-dependent siRNA-mediated inhibition of OP18 gene expression showed strong effects in the G3 model cell line ECV-304, but only moderate down-regulation of OP18 to levels of 31% at 24 h after transfection in RT-4 (Fig. 2 A).

**Fig 2.**
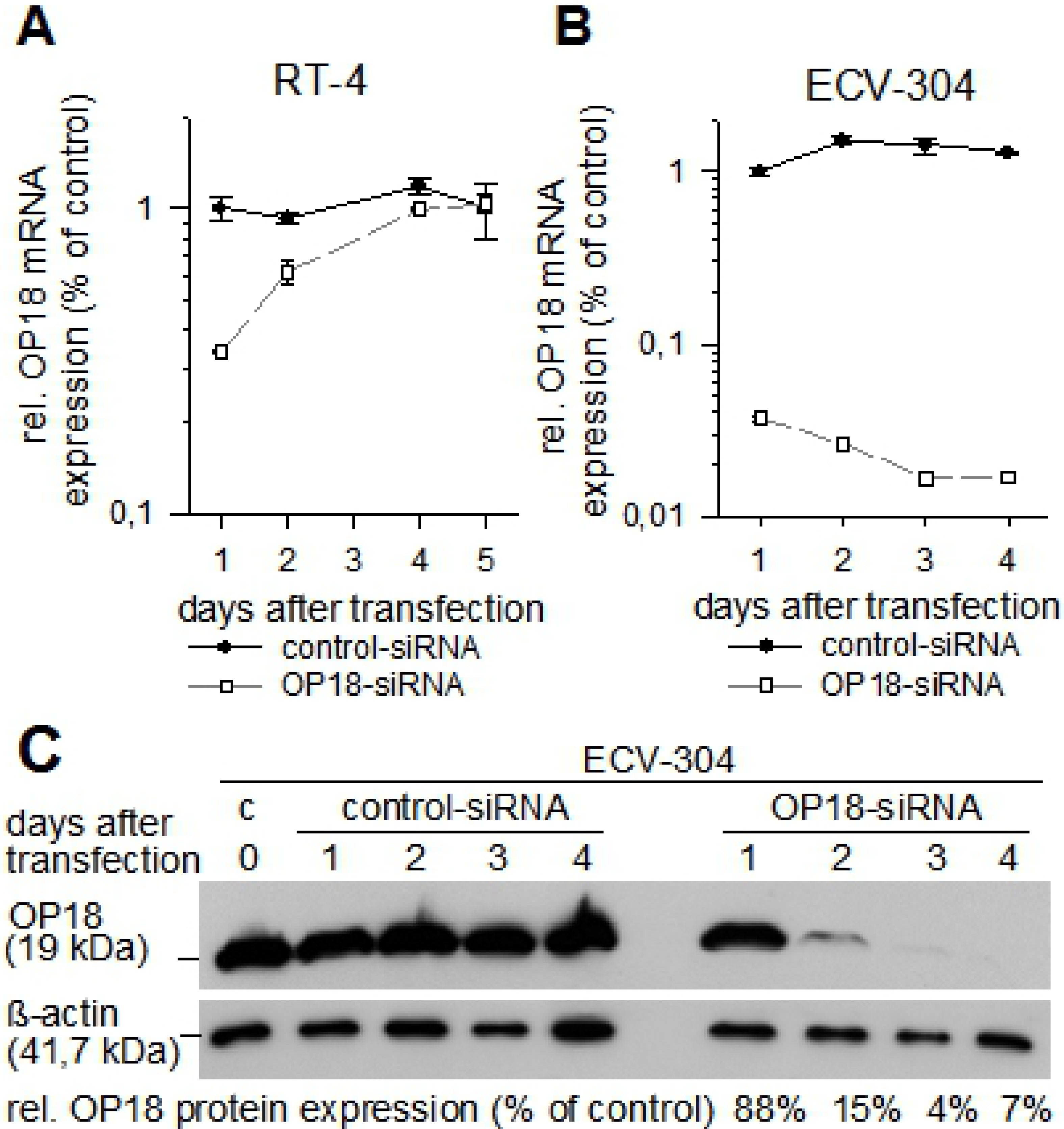
Validation of siRNAs. The two cell lines RT-4 **(A)** and ECV-304 **(B)** were transfected with OP18-directed siRNA or control siRNA at 30 nM. Total RNA was prepared after transfection at the indicated time points and levels of OP18- and RPLP0-mRNA were detected by qRT-PCR in triplicate. Symbols represent mean relative OP18 mRNA expression ± standard deviation as normalized to RPLP0. **(C)**, relative OP18 protein amount in siRNA-treated ECV-304 cells. Cells were transfected with control-siRNA or OP18-siRNA (each at 30 nM). At 0, 1, 2, 3, and 4 days after transfection, cells were lysed, and OP18 protein and beta-actin were detected using western blotting. Signals of OP18 were normalized to beta-actin

Percentages of OP18 protein suppression in comparison to control cells is indicated in the lower panel of the blot.

Thereafter, OP18 mRNA expression increased to the level detected in control-siRNA treated RT-4 cells. In contrast, in ECV-304 cells, the OP18 expression was inhibited efficiently to levels of approximately 2% of relative expression (Fig. 2 B). At the OP18 protein level, suppression was investigated for ECV-304 only (Fig. 2 C). A substantial decrease of OP18 protein signal was observed at days 2, 3, and 4 after transfection which relates to 15%, 4%, and 7% suppression, respectively (as compared to control siRNA).

### 3.3 Suppression of OP18 is associated with reduced cell proliferation in ECV-304

Next, we investigated a possible correlation of siRNA-mediated suppression of OP18 and cell proliferation in RT-4 and ECV-304 (Fig. 3).

**Fig 3.**
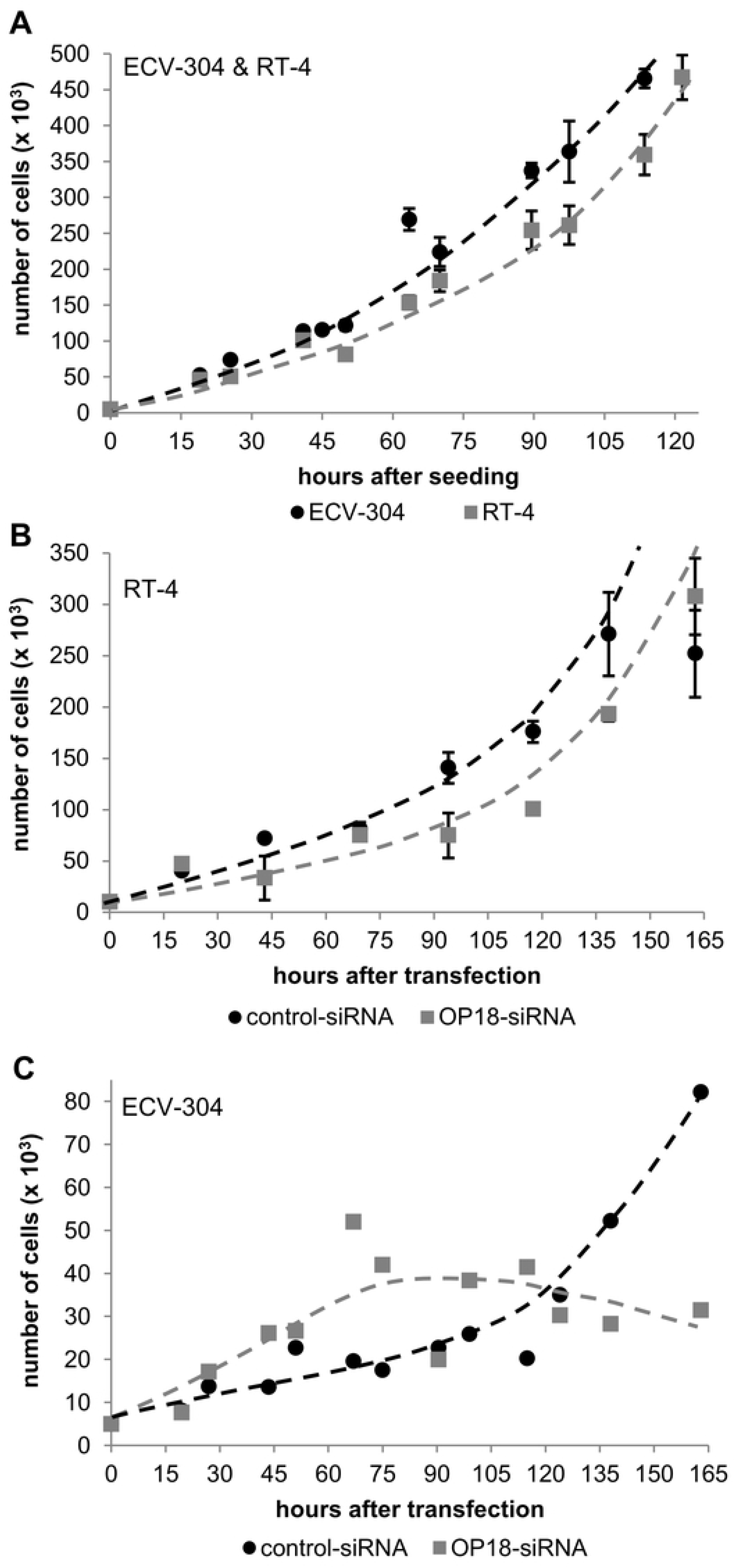
Phenotypic characteristics of OP18-suppressed BCa cell lines. Proliferation rates of untreated RT-4 and ECV-304 **(A)** siRNA-treated RT-4 **(B)** and siRNA-treated ECV-304 **(C)**. BCa cell lines were transfected in duplicate with 30 nM of OP18-siRNA or control-siRNA. Numbers of viable cells at the indicated time points after transfection were determined by trypan blue staining. The data indicate mean values of 3 experiments ± standard deviation.

Analysis of cell growth of untreated and siRNA-treated RT-4 and ECV-304 cells was conducted. Untreated RT-4 cells had slightly longer doubling times by a factor of approximately 1.5 compared to ECV-304 (Fig. 3A). After transfection with functional or control siRNA, RT-4 still displayed nearly similar proliferation rates (Fig. 3B). Conversely, the proliferation of ECV-304 transfected with OP18-directed siRNA showed substantially decreased cell proliferation after day 4, when compared to treatment with control siRNA (Fig. 3C).

### 3.4 OP18-suppressed ECV-304 cells undergo apoptosis

To study the potential relationship between OP18 suppression and apoptosis, we analyzed the expression of apoptosis-related genes, (pro-apoptotic: BAX and CC3; anti-apoptotic: TC3) after transfection of ECV-304 cells with OP18-directed siRNA. While the expression level of BAX mRNA did not differ significantly between OP18-suppressed cells and controls (Fig. 4 A), the pro-apoptotic mRNA ratio CC3:TC3 increased progressively at day 4 after the suppression of OP18 (Fig. 4 B).

**Fig 4.**
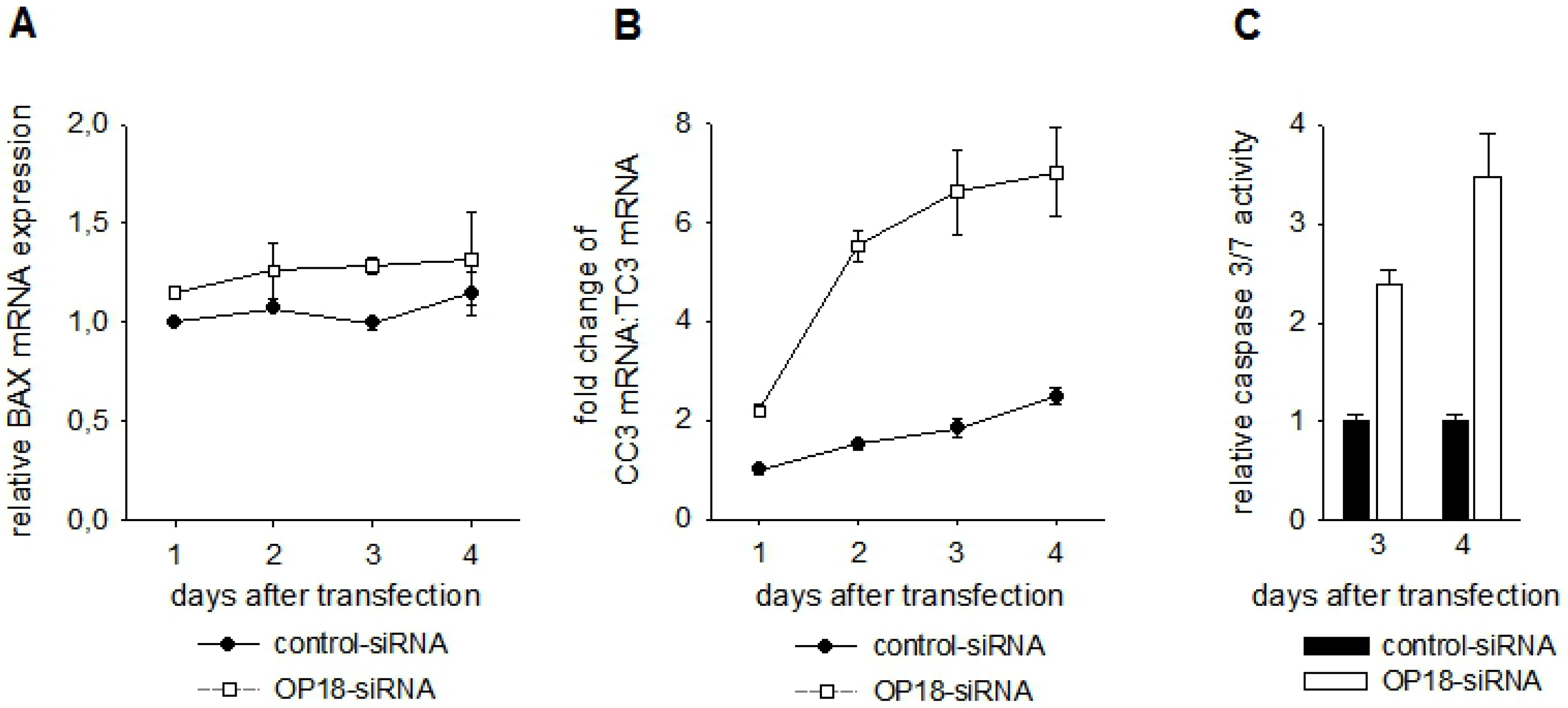
Apoptotic effects of OP18-suppression in ECV-304. Relative expression of the pro-apoptotic BAX mRNA **(A)** and increase (‘fold change’) of the pro-apoptotic mRNA-ratio CC3:TC3 **(B)**. ECV-304 cells were transfected in duplicate with 30 nM OP18- and control-siRNA and total RNA was prepared at 1, 2, 3, 4 days after transfection. Levels of BAX, CC3, TC3 and RPLP0 mRNA were detected in triplicate by qRT-PCR. Data indicate mean values ± standard deviation and are representative of three independent experiments. **(C)**, induction of caspase 3/7 activity after suppression of OP18. ECV-304 cells were transfected in triplicate with OP18- and control-siRNA (each 30 nM) and the relative caspase 3/7 activity as normalized to cell viability was quantified at day 3 and 4 after transfection. Data indicate mean values ± standard deviation and are representative of three independent experiments.

To further investigate the induction of apoptosis in OP18-suppressed ECV-304, relative caspase 3/7 activity was determined at the protein level (Fig. 4 C). At day 3 and 4 after transfection, caspase 3/7 activity was enhanced by 2.5- and 3.5-fold, respectively.

### 3.5 Suppression of OP18 increases chemo-sensitivity in ECV-304

In a more therapeutically oriented perspective, the role of OP18 in sensitivity of ECV-304 for the chemotherapeutic agent cisplatin was studied (Fig. 5).

**Fig 5.**
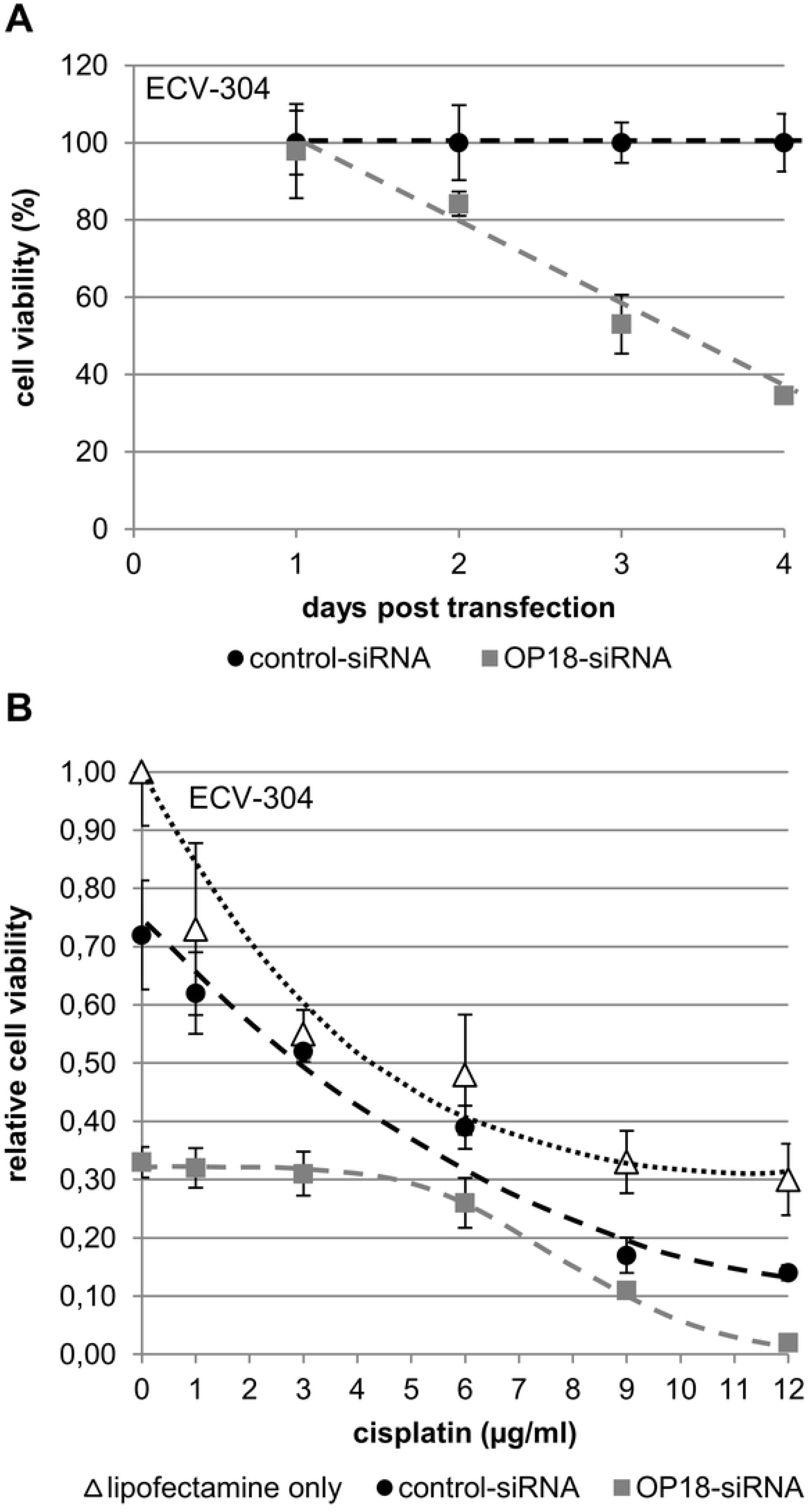
Chemo-sensitivity of BCa cells after suppression of OP18. ECV-304 cells were transfected in triplicate with each 30 nM of OP18- and control siRNA. **(A)**, cell viability of siRNA-treated ECV-304 cells. The viability of ECV-304 cells was quantified with the colorimetric MTS-assay at 1, 2, 3, 4 days after transfection in triplicate (30 nM OP18- and control-siRNA). Symbols indicate mean values ± standard deviation of three independent experiments. **(B)** Cells were treated for 24 h with different concentrations of cisplatin (0, 1, 3, 6, 9, and 12 μg/ml) 48 h after transfection followed by quantification of cell viability. Data indicate mean values ± standard deviation and are representative of three independent experiments.

Analysis of cell viability of OP18-suppressed ECV-304 cells showed a decrease of 35% at day 4 after transfection with functional siRNA when compared to cells transfected with control siRNA (Fig. 5B).

Subsequently, ECV-304 cells were exposed to varying concentrations of cisplatin for 24 h at day 2 post-transfection with siRNA. Notably, treatment with cisplatin had an additive effect on loss in relative viability in ECV-304: At high concentrations of cisplatin (9 and 12 μg/ml), a progressively severe effect was observed on cell viability of OP18-suppressed cells as compared to controls. At a cisplatin concentration of 12 μg/ml, the decrease in cell viability of OP18-suppressed ECV-304 cells was in the order of one magnitude while in the absence of cisplatin, this was only approximately twofold (Fig. 3C and Fig. 5, “0” cisplatin). At low concentrations of cisplatin (1, 3 and 6 μg/ml), the effect on relative cell viability was similar in OP18-suppressed cells and controls.

## 4 DISCUSSION

### 4.1 This study provides functional insights into the biological role of OP18 and its involvement in malignant cell proliferation

Prognostic value has been assigned to OP18 in different tumor entities as based on tissue biopsies [17–20]. Recently, Hemdan *et al.* investigated the role of OP18 in BCa [21]. In line with the present study, they used siRNAs, but a commercial set of siRNAs, not overlapping with the designed siRNAs used in this study for suppression of OP18 mRNA, supporting our findings.

In contrast to Hemdan *et al.*, we used liquid biopsies, two cell culture models and molecular analyzes rather than tissues and histopathological staining. The use of whole urine samples is advantageous because urine is readily available and can be collected in high frequency, e.g. to monitor patients with BCa with a relatively high recurrence rate. Moreover, molecular markers can be detected in a standardized manner by qPCR, whereas histopathological examinations of tumor tissue strongly depend on the pathologist or pattern recognition imaging software.

### 4.2 OP18 as potential therapeutic bladder cancer target and tumor marker

The suppression of OP18 by siRNA was observed on the levels of mRNA and protein which suggests to further test a wide repertoire of inhibitors to therapeutically address more advanced tumor stages of bladder cancer to less malignant stages. We anticipate that instillation of drugs into the bladder produces a scenario of drug application that is closer to a local rather than a systemic application which implies several fundamental advantages such as increased local concentration, i.e., increased delivery to tumor cells, higher stability, and decreased side effects.

This study strongly suggests OP18 to be a molecular marker and a cause for the disease. We assume that OP18-specific RNA contained at elevated amounts in the urine of BCa patients at advanced tumor stages reflects increased OP18 expression levels in tumor tissue. It is reasonable that OP18 serves as a tumor marker closely related to malign molecular events, rather than indirectly reflects a tumorigenic cellular process. Thereby, OP18-specific RNA as a marker directly monitors the disease. Further, to improve sensitivity and specificity of this marker, combinations of markers in the line of OP18-specific RNA might even give rise to substantially improved diagnostics of BCa and in a non-invasive setting. For example, the analysis of urinary mRNAs in this study revealed an improved relationship between the mRNA-ratio OP18/UPK1A and poorly differentiated (G3) and muscle-invasive (≥ pT2) BCa cancer states. Our data strongly indicate that this strategy has great potential for future accurate and non-invasive diagnostic developments in case of bladder cancer and beyond.

## 5 CONCLUSIONS

OP18 expression seems to be necessary for malignant cell proliferation in human cells derived from bladder cancer. In a diagnostic perspective, urine RNA levels (OP18:UPK1A) serve as a molecular marker for the invasive disease. In mechanistic terms, over-expression of OP18 seems to be necessary for maintaining the malignant state of BCa cells as its suppression results in an increased chemo-sensitivity and apoptosis. Hence, the gene expression of OP18 represents a rational therapeutic target and diagnostic readout.

## 6 ACKNOWLEDGEMENTS

We thank Drs. C. Höppner, M. Horn, T. Menke, S. Thomas, and J. Träder for collecting samples, B. Thode and B. Branke for technical assistance.

## 8 SUPPORTING INFORMATION CAPTIONS

S1 Table: Sequences and characteristics of qPCR amplicons

S2 Table: Most effective siRNA sequence and scrambled RNA sequence

